# Towards building a World Model to simulate perturbation-induced cellular dynamics by AlphaCell

**DOI:** 10.64898/2026.03.02.709176

**Authors:** Guohui Chuai, Xiaohan Chen, Xingbo Yang, Cheng Zhang, Kairu Qu, Yiheng Wang, Wannian Li, Jingya Yang, Duanmiao Si, Feiyang Xing, Yicheng Gao, Siqi Wu, Shaliu Fu, Bing He, Qi Liu

**Affiliations:** Bioinformatics Department, School of Life Sciences and Technology, Tongji University, Shanghai, 200092, China; National Key Laboratory of Autonomous Intelligent Unmanned Systems, Frontiers Science Center for Intelligent Autonomous Systems, Ministry of Education, Shanghai Key Laboratory of Intelligent Autonomous Systems 201804, China; Shanghai Qi Zhi Institute, 200232, China

## Abstract

Predicting cellular responses to perturbations is crucial for therapeutic discovery, yet experimental screening is severely constrained by the combinatorial vastness of biological space. While computational simulations offer a scalable alternative, current models are limited by the incomplete latent representation——mainly relying on highly variable genes in feature representation; the poor genome-wide reconstruction fidelity; and the ungeneralizable dynamic laws across diverse contexts. Consequently, they fail to mechanistically transfer learned dynamics to unseen cellular contexts. To address these systemic flaws, we introduce AlphaCell, a generative Virtual Cell World Model that unifies genome-wise representation with continuous state transition modeling. AlphaCell achieves three synergistic innovations: (1) Latent Manifold Rectification, processing the full protein-coding transcriptome to construct a differentiable Virtual Cell Space, effectively filtering noise while preserving intrinsic cellular topology; (2) Biological Reality Reconstruction, utilizing a massive, knowledge-rich decoder to translate abstract latent states back into high-fidelity, genome-wide expression profiles; and (3) Universal State Transition, applying Optimal Transport Conditional Flow Matching to model perturbations as continuous, deterministic vector fields. By abstracting perturbation mechanisms into generalized dynamic laws, AlphaCell makes robust prediction of perturbation responses in a compositional generalization scenario and enables zero-shot prediction of cellular dynamics in entirely unseen cellular contexts, providing a foundational engine for cellular-context-generalizable perturbation prediction and perturbation-induced cellular dynamics simulation.

## Introduction

Predicting how cells respond to perturbations is central to therapeutic discovery^1,2^. While advances in single-cell genomics enable large-scale screening across specific contexts^3,4^, experimental approaches remain fundamentally constrained by the combinatorial vastness of biological space. It is cost-prohibitive and labor-intensive to empirically test every potential genetic or chemical intervention across every cellular context. To overcome these limitations, computational biology must evolve from descriptive inference to predictive simulation—generating AI Virtual Cells (AIVC) as digital twins of living cellular systems to simulate and predict the cellular dynamics^5^. These AI-driven virtual models serve as computational surrogates for physical experiments, leveraging vast biological datasets to mathematically encode cellular states and dynamic rules^6^. By capturing the intrinsic logic of how cells maintain identity and respond to the perturbations, AIVCs offer a scalable platform to test biological hypotheses in silico, circumventing the bottlenecks of traditional experimentation.

From a broad perspective, cellular perturbation prediction is a kind of cellular state transition simulations, representing one specific scenario within the cellular dynamic modeling. Despite the urgent need for such simulation capabilities, current computational frameworks related to AIVC remain fragmented across distinct technical routes, each relying on simplifying abstractions to render the problem tractable. A first paradigm focuses on latent arithmetic, where models like scGen^7^, CPA^8^, and biolord^9^ utilize variational autoencoders to disentangle cellular signals from technical covariates, explicitly modeling perturbation responses as linear vector shifts added to a latent representation derived from highly variable genes (Fig. 1b). A second paradigm incorporates mechanistic constraints, exemplified by GEARS^10^ and CASCADE^11^, which constrain the solution space using prior biological knowledge graphs or causal inference frameworks to guide rational cellular state transition predictions (Fig. 1c). A third paradigm targets population dynamics: CellOT^12^ leverages optimal transport theory to learn discrete mappings between unpaired distributions, while CellFlow^13^ adopts flow matching to model continuous cellular trajectories, albeit typically restricted to low-dimensional principal component spaces (PCA) (Fig. 1d). Finally, foundation models like scGPT^14^ and STATE^15^ leverage massive datasets via transformer architectures for cellular representation and downstream perturbation predictions; specifically, scGPT applies masked modeling to restricted gene tokens, while STATE utilizes set-based attention and maximum mean discrepancy (MMD) loss to align population statistics (Fig. 1e). However, regardless of the methodology advances, these approaches converge on three fundamental architectural flaws that hinder the development of high-fidelity virtual cells in the simulations of cellular dynamics:

(1) Latent representation incompletion: nearly all models, including recent flow-based and foundation approaches, rely on heuristic feature selection, typically restricting inputs to the top 1,000 ∼ 2,000 highly variable genes (HVGs) or a fixed gene set (for example, the L1000 gene set) as a feature representation of the virtual cell as a feature representation of the virtual cell^16^. This input truncation violates the theoretical completeness required to rigorously define a cellular state. Since it systematically excludes low-abundance but high-information regulatory drivers (e.g., master transcription factors and receptors), the resulting feature space is fundamentally incomplete and heavily biased toward the training distribution. Furthermore, raw single-cell sequencing data, which is sparse, discrete, and noisy, is inherently ill-suited for defining continuous dynamics^17^. Traditional latent spaces often passively inherit these topological defects rather than rectifying them, failing to provide a mathematically valid workspace for physical simulation of the cellular dynamics.
(2) Biological reconstruction distortion: existing latent models often lack a robust mechanism to translate abstract latent predictions back into high-fidelity biological reality. Without a powerful genome-wide decoder to ensure reconstruction fidelity across the whole genome, mathematical operations within the latent space may generate biological hallucinations that cannot be grounded in measurable transcriptomic phenotypes.
(3) Dynamic transferability deficiency: existing methods generally model perturbation as a discrete jump or model continuous flows directly within restricted HVG spaces. Lacking a universal latent cell embedding that unifies diverse datasets, these approaches fail to learn generalizable laws of cellular state transitions that apply across diverse cellular contexts.

**Figure 1.**
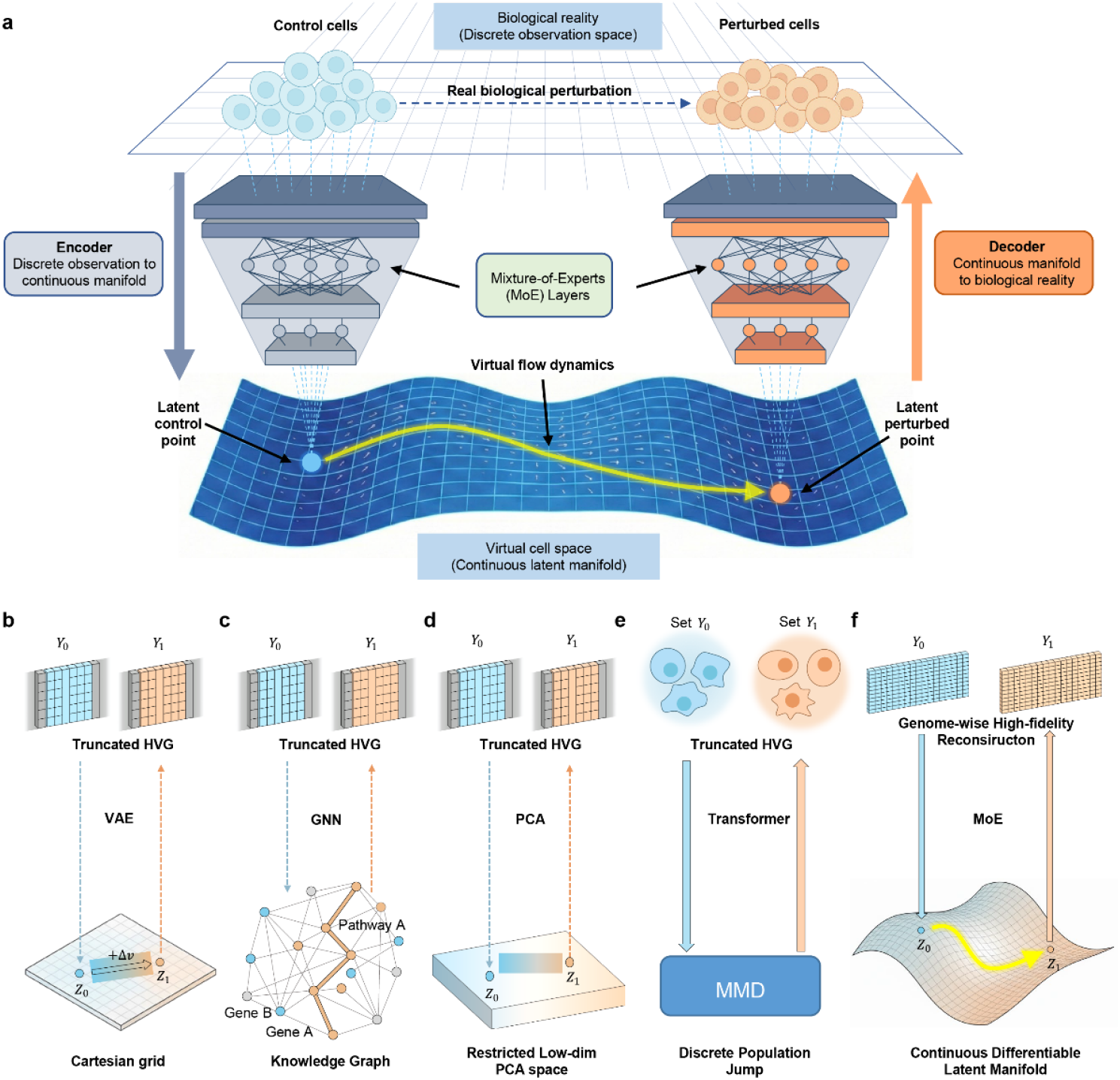
The AlphaCell Virtual Cell World Model framework and various cellular state transition paradigms. (a) The overarching architecture of AlphaCell. (b–f) Conceptual comparison between existing computational paradigms and AlphaCell. b, Latent arithmetic models (e.g., VAE-based) assume rigid linear vector shifts in flat Cartesian grids based on truncated highly variable genes (HVGs). c, Mechanistic constraint models (e.g., GNN-based) rely on discrete state transitions guided by prior knowledge graphs. d, Restricted subspace flow models capture continuous trajectories but are confined to narrow, low-dimensional spaces (e.g., PCA) using truncated gene inputs. e, Set-based foundation mapping utilizes Transformers and Maximum Mean Discrepancy (MMD) to align population statistics, resulting in discrete population jumps without individual continuous dynamics. f, AlphaCell establishes a complete Virtual Cell World Model: it processes genome-wise profiles without truncation, constructing a continuous differentiable latent manifold where perturbation responses are modeled as continuous physical flows and reconstructed with high fidelity.

Consequently, these structural deficits culminate in a failure of generalization: tethered to the superficial statistics of their training distributions, current models are rendered incapable of mechanistically transferring learned dynamics to entirely unseen cellular contexts (zero-shot prediction). To addressing these systemic limitations, we proposed that a paradigm shift from fragmented analytical tools toward a unified ***World Model*** is needed. Originating from embodied artificial intelligence, a ***World Model*** is defined as an internally learned representation of an environment capable of simulating future states and governing dynamic transitions^18,19^. Accordingly, to construct a **Virtual Cell World Model (Figure 1a,f)**, the system must be an integrated framework satisfying three indispensable and interdependent requirements: (1) a genome-wise representation that rectifies discrete observations into a computable, continuous latent manifold; (2) a high-fidelity observation interface that faithfully translates abstract latent states back into biological reality with minimum distortion; and (3) a transferable dynamic mechanism to simulate continuous cellular transitions across diverse contexts. To this end, we introduce AlphaCell, a generative **Virtual Cell World Model** specifically designed to fulfill these criteria. Briefly, AlphaCell fundamentally reimagines cellular dynamic modeling by establishing a rigorous **Virtual Cell Space**—a low-dimensional latent manifold where the digital twin of a cell resides, being observed, and evolves. To realize this vision, we implemented AlphaCell through three synergistic theoretical innovations:

(1) Latent Manifold Rectification: to resolve the latent representation incompletion, the AlphaCell ***Base Model Encoder*** functions as a Virtual Cell Space builder. By processing the full protein-coding transcriptome rather than restricted gene sets, it projects complex and genome-wise biological states into a robust cell state embedding within a continuous latent space (Virtual Cell Space), which transforms discrete, noisy transcriptome observations into a differentiable mathematical substrate. It creates a robust information bottleneck that filters technical noise while preserving intrinsic cell state topology, providing the necessary foundational workspace where cell states become computable.
(2) Biological Reality Reconstruction: to bridge the Virtual Cell Space with biological reality and solve the biological reconstruction distortion, AlphaCell incorporates a massive, knowledge-rich ***Base Model Decoder*** as a precise observation interface. This component ensures that any given cell state embedding in the Virtual Cell Space can be translated back into a biologically valid, genome-wise expression profile. This guarantees that cell state embeddings retain strict biological fidelity, preventing hallucinations and ensuring simulated trajectories can be reliably decoded into original phenotypic reality.
(3) Universal State Transition: to resolve the dynamic transferability deficiency, the AlphaCell ***Flow Model*** acts as the physics engine within this initialized space. Leveraging Optimal Transport Conditional Flow Matching^20,21^, it transforms perturbations from discrete jumps into continuous flows. By learning deterministic vector fields that transport cell state embeddings from control to perturbed states, AlphaCell mathematically models the non-linear evolution of cell states, where perturbation responses are computed as precise physical flows acting in the Virtual Cell Space. Therefore, AlphaCell is able to effectively abstract perturbation mechanisms into generalized dynamic laws that are transferable across specific cell identities, enabling the mechanistic transfer of learned dynamics to entirely unseen cellular contexts.

## Result

### The AlphaCell framework: a Virtual Cell World Model

We constructed AlphaCell as a unified generative system designed to simulate a **Virtual Cell World Model**—a digital twin capable of simulating and predicting cellular dynamic (Fig. 1a).

As a World Model, the AlphaCell architecture is organized into a synergistic triad: (1) a Virtual Cell Space builder (***the Base Model Encoder***) that compresses genome-wise cellular context inputs into a computable latent manifold; (2) a high-fidelity Observation Interface (***the Base Model Decoder***) that translates abstract latent states back into measurable biological reality with minimum distortion; and (3) a Dynamic Physics Engine (***the Flow Model***) that dictates the continuous laws of cellular state transition within the Virtual Cell Space.

AlphaCell realizes these capabilities through a multi-stage architecture trained on an unprecedented scale. The ***Base Model*** (comprising the Encoder and Decoder) was trained on over 220 million single cells (140 million observational transcriptomes from CZ CELLxGENE^22^ with 80 million profiles from the Tahoe dataset^23^) to construct the Virtual Cell Space and Observation Interface. Subsequently, the ***Flow Model*** was trained on 90 million perturbed profiles (80 million profiles from the Tahoe dataset and nearly 10 million profiles from pharmacological, e.g. Sciplex^24^, and genetic overexpression screens^1^) to define the generalized dynamic laws of cellular evolution.

#### Constructing a continuous Virtual Cell Space via genome-wise distillation and manifold rectification

The primary foundation of the AlphaCell framework is the construction of a rigorous Virtual Cell Space—a continuous manifold where the digital twin of any cell resides. To ensure this space serves as a valid mathematical substrate for continuous physical simulation, it was engineered to satisfy four rigorous design criteria: (1) Informational Completeness, incorporating genome-wise cellular context inputs rather than truncated HVG feature sets; (2) Manifold Differentiability, transforming discrete, sparse observations into a dense continuous space to support dynamic trajectories; (3) Batch Invariance, establishing a universal latent manifold independent of technical artifacts; and (4) Semantic Fidelity, ensuring the abstract representation retains full biological information required for precise decoding.

To achieve the first criterion, Informational Completeness, we prioritized rigor at the input stage. AlphaCell processes Unique Molecular Identifier (UMI)-based sequencing data, standardized to log(1+CP10k) and discretized via a fine-grained 100× tokenization strategy to capture subtle gene dosage variations often lost in standard normalization. Unlike prevailing methods that restrict inputs to ∼2,000 HVGs, AlphaCell processes the full set of 19,253 HGNC protein-coding genes^25^. We specifically aligned data to the HGNC standard to eliminate semantic ambiguity often found in raw genomic annotations (e.g., GENCODE^26^), where a single gene symbol may map to multiple Ensembl IDs^27^. This ensures a strict bijective mapping between biological entities and model inputs, fulfilling the theoretical informational completeness required to rigorously define a cellular state.

To achieve the second criterion, Manifold Differentiability, we designed a Mamba-Transformer hybrid encoder to transform the high-dimensional, sparse, and noisy input into a tractable Virtual Cell Space. Crucially, this encoder is not merely a dimensionality compressor, but a manifold rectifier. By combining the linear scaling of State Space Models (Mamba)^28^ with the global attention of Transformers^29^, the encoder captures long-range regulatory dependencies across the entire feature space. The encoder projects the cellular state into a continuous latent space organized as 32 coupled state channels (dimensions 32×128) (Fig. 2a). This structural information bottleneck filters stochastic dropout and technical noise. Furthermore, to explicitly prevent structural fragmentation and enforce the topological smoothness of this continuous space, we applied stringent L2 regularization to the latent representations during the Fine-tuning stage. Together, the architectural bottleneck and L2 constraints ensure the manifold remains dense and mathematically differentiable, primed for continuous dynamic simulation.

**Figure 2.**
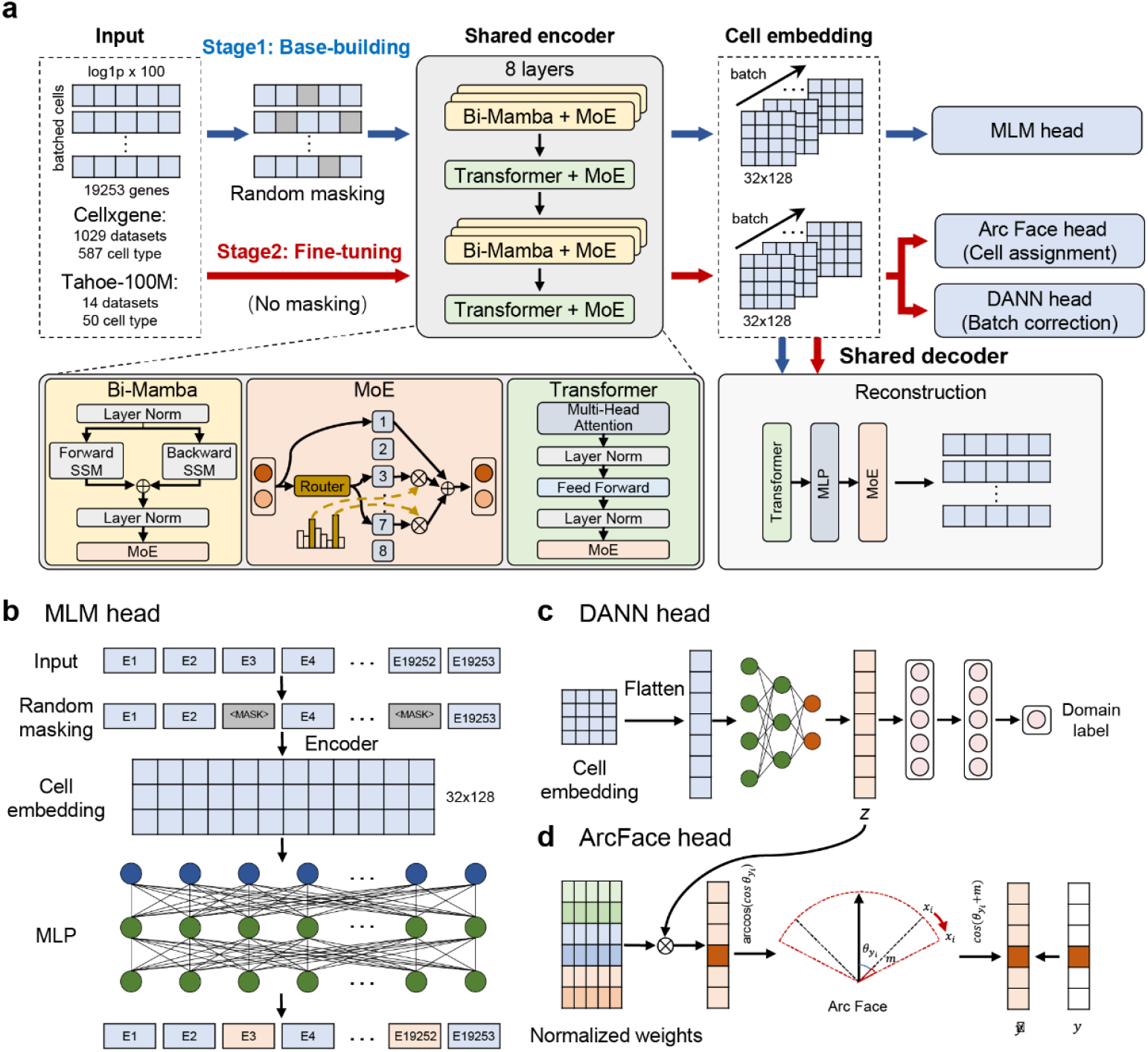
Architecture and training paradigm of the AlphaCell Base Model for Virtual Cell Space construction. (a), Schematic of the AlphaCell Base Model architecture and training curriculum. Training spans two phases: Stage 1 (Base-building) utilizing masked self-supervision, and Stage 2 (Fine-tuning) incorporating ArcFace and DANN constraints, with a Shared MoE Decoder ensuring high-fidelity whole-genome reconstruction. (b), The Masked Language Modeling (MLM) head used in Stage 1. By reconstructing randomly masked gene tokens, the model learns the fundamental syntax of gene co-expression without supervision. (c), The Domain Adversarial Neural Network (DANN) head used in Stage 2. It applies a gradient reversal layer to the flattened cell embedding, adversarially filtering out technical batch signatures to ensure the universality of the Virtual Cell Space. (d), The ArcFace head used in Stage 2. It imposes angular margins on normalized weights to maximize the separability of distinct biological states, refining the topological structure of the latent manifold.

To achieve the third criterion, Batch Invariance, we employed a two-stage learning curriculum (Fig. 2a). While the initial Base-building stage constructs the foundational topology, we integrated a channel-wise Domain Adversarial Neural Network (DANN)^30^ (Fig. 2c) during the Fine-tuning stage. By adversarially stripping technical signatures from each state channel individually, AlphaCell establishes a unified latent manifold where the position of a cell is determined solely by biological identity, strictly disentangled from batch artifacts.

Finally, to achieve the fourth criterion, Semantic Fidelity, which ensures the abstract manifold retains full and accurate biological information, we designed a two-stage, reconstruction-driven curriculum. During the Base-building stage, the model processes randomly masked inputs and is optimized via a joint Masked Language Modeling (MLM)^31^ and reconstruction objective (Fig. 2b). Because the input is heavily corrupted, this serves as a rigorous denoising reconstruction task, compelling the encoder to internalize the fundamental syntax of gene co-expression to recover the missing biological signals. Subsequently, during the Fine-tuning stage, the model transitions to processing unmasked, full-context inputs. In this phase, we integrated an ArcFace head^32^ (Fig. 2d) to perform explicit cell assignment and clustering, imposing angular margins to maximize the separability of distinct biological identities. Moreover, a concurrent unmasked reconstruction objective is maintained alongside ArcFace and the abovementioned DANN task. This ensures that the aggressive structural shaping and batch-correction do not collapse essential transcriptomic details, thereby locking in the strict biological fidelity of the latent representations. Ultimately, validating this semantic fidelity requires proving that these abstract cell embeddings can be seamlessly translated back into genome-wise phenotypes, necessitating a uniquely powerful observation interface.

Through this rectification process, AlphaCell elevates fragmented and noisy transcriptomic snapshots into a continuous, universally aligned Virtual Cell Space. This rectified manifold not only preserves the intrinsic biological topology of the digital twins, but also more importantly, provides the exact differentiable substrate required to simulate the non-linear trajectories of cellular evolution.

#### Reconstructing biological reality from Virtual Cell Space via an inverted pyramid Mixture-of-Experts architecture

A rigorous Virtual Cell Space is only scientifically valuable if its abstract coordinates can be translated back into biological reality. To bridge the cell state embedding back to the observable transcriptome, we implemented an “Inverted Pyramid” architecture. Unlike typical autoencoders where the encoder and decoder are roughly symmetric, AlphaCell attaches a massive 1.2-billion-parameter Mixture-of-Experts (MoE)^33,34^ Decoder directly to the latent manifold (Fig. 2a).

This structural asymmetry is deliberate. The Decoder functions as a deep biological knowledge base, encoding the complex, non-linear gene co-expression priors required to expand the compacted cell state embedding back into the full 19,253-gene profile. By leveraging the sparse activation of MoE, the model scales its parameter capacity to memorize vast regulatory dependencies without incurring prohibitive computational costs. This design ensures that despite the high compression of the Virtual Cell Space, any valid cell state embedding within the manifold can be translated into a biologically accurate genome-wise expression profile with high precision (ROC-AUC > 0.96, Pearson > 0.7, MAE < 0.25). This rigorous reconstruction capability validates that the cell state embedding within the Virtual Cell Space successfully captures the comprehensive state of the cell, providing a reliable ground truth for downstream cellular dynamic simulation.

#### Simulating cellular state transition via optimal transport conditional flow matching

With the static Virtual Cell Space established, the AlphaCell ***Flow Model*** functions as the system’s physics engine, defining how biological states evolve under external conditions. To enable such simulation, the latent manifold must support continuous temporal evolution rather than discrete jumps. However, a fundamental challenge in training such a dynamic model is the destructive nature of single-cell sequencing, which precludes observing the temporal trajectory of a single cell. To overcome this, we leverage Optimal Transport Conditional Flow Matching (OT-CFM)^20,21^ (Fig. 3b) coupled with a dynamic intra-batch Optimal Transport strategy (Fig. 3c). During training, control and perturbed populations are sampled and matched on-the-fly within each mini-batch based on their proximity within the Virtual Cell Space. This process constructs probabilistic geodesic paths from unpaired population snapshots, allowing the model to learn a deterministic vector field *v*(*z,t,c*) that transports a cell state embedding from its control state (*z*_ctrl_) to its perturbed state (*z*_pert_) along the optimal trajectory in the Virtual Cell Space.

**Figure 3.**
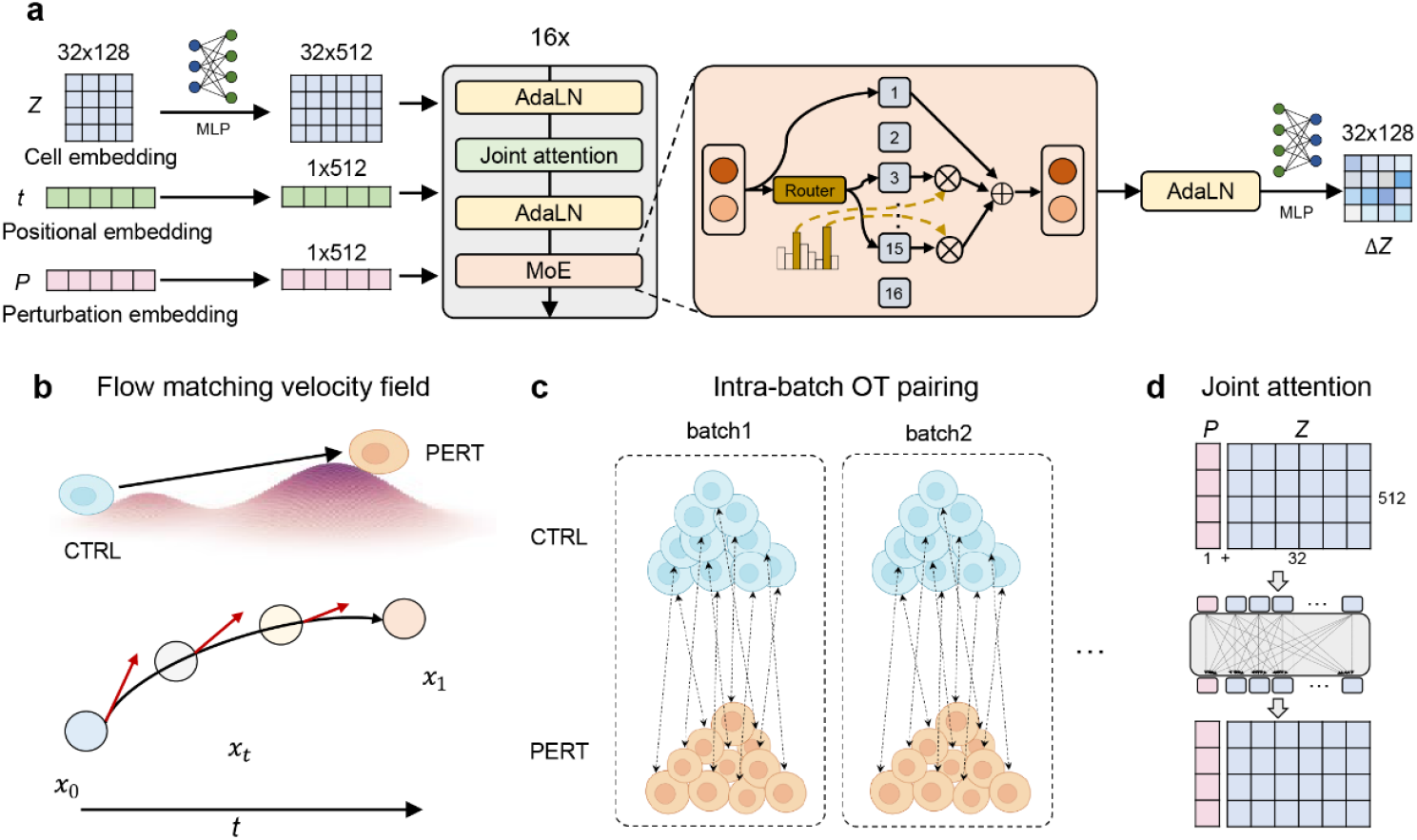
Generative dynamics with MoE-based Optimal Transport Conditional Flow Matching. (a), Architecture of the AlphaCell Flow Model. Inputs, including cell state, time, and perturbation, are processed through a backbone featuring Adaptive Layer Normalization (AdaLN), Joint Attention, and a Shared/Routed Mixture-of-Experts (MoE) to dynamically encode perturbation-specific dynamics. (b), Optimal Transport Conditional Flow Matching (OT-CFM). The model learns a deterministic vector field transporting control states (blue) to perturbed states (orange) along continuous geodesic trajectories. (c), Dynamic Intra-batch Optimal Transport. To handle unpaired data, control and perturbed populations are matched on-the-fly within mini-batches to define optimal training paths. (d), Joint Attention. This mechanism modulates cell state channels based on perturbation semantics.

This continuous formulation allows AlphaCell to simulate the non-linear dynamics of biological transitions. To further handle the immense complexity of diverse perturbation mechanisms, the Flow Model employs an advanced Shared and Routed Mixture-of-Experts (MoE) architecture (Fig. 3a). To ensure precise control over the trajectory, perturbation signals are injected via Adaptive Layer Normalization (AdaLN)^35^ and a Joint Attention mechanism (Fig. 3d), which dynamically modulates the interaction between the perturbation embedding and the cell state. Instead of forcing a single dense network to average across potentially conflicting dynamics, this design significantly expands the model’s capacity to encode a vast repertoire of vector fields. By utilizing conditional computation, the model effectively mitigates the gradient conflicts that typically arise when learning heterogeneous perturbation effects simultaneously, preventing catastrophic interference between varying conditions. Crucially, because these laws of motion are learned on the universal Virtual Cell Space rather than on dataset-specific features, AlphaCell achieves mechanistic transferrable ability cross diverse cellular context. A clearly depiction and comparison of this point between AlphaCell and existing models are shown in Figure 1b-e, as described aforementioned. Once the model masters the vector field induced by a specific perturbation on common cell types, it can mathematically apply this force to the cell state embedding of entirely unseen cell types, enabling the prediction of perturbation dynamics in biological contexts that were physically untested.

### Robust prediction of perturbation responses in a compositional generalization scenario

Cellular perturbation response prediction is a kind of cellular state transition simulation. In this study, we first evaluated the generalizable capacity of cellular state transition modeling of AlphaCell on a critical perturbation response prediction task, i.e, the compositional perturbation response prediction generalization task—predicting the response of a known cell type to a known perturbation in a novel “cell-perturbation” pairing configuration. This setup evaluates the model’s first level of cellular context generalization ability in the cellular state transition modeling: testing whether the perturbation mechanism has been successfully abstracted into a generalized dynamic law capable of reliably operating upon the distinct initial state of a newly paired cellular identity. We benchmarked AlphaCell against a comprehensive suite of baselines, including linear models, specialized predictors (CPA, GEARS, CASCADE), and recent foundation models (scGPT, STATE), across three diverse datasets spanning genetic and chemical modalities: OTF, Sciplex, and Tahoe-100M. Crucially, this benchmark represents an asymmetrical challenge favoring the baselines. All competing models were firstly restricted to the top 2,000 highly variable genes (HVGs)—a standard practice to mitigate sparsity that theoretically caps the model’s ability to capture complex regulatory logic. Then, to ensure a comprehensive comparison, the competing models (if applicable) were also evaluated using the full 19,253-geneset. We observed that directly extending these models from HVGs to the full gene set led to a significant degradation in their predictive performance. This deterioration exposes a fundamental structural vulnerability in existing architectures: without a robust manifold rectification mechanism, these models succumb to the curse of dimensionality and are overwhelmed by the zero-inflated noise inherent in genome-scale observations. In contrast, AlphaCell maintained a high performance on the genome-wise task, consistently surpassing all baselines across all metrics (Fig. 4). This resilience demonstrates that AlphaCell’s architecture, specifically its continuous latent manifold and high-fidelity decoder, effectively captures complex regulatory logic at the whole-genome scale without being compromised by pervasive sparsity.

**Figure 4.**
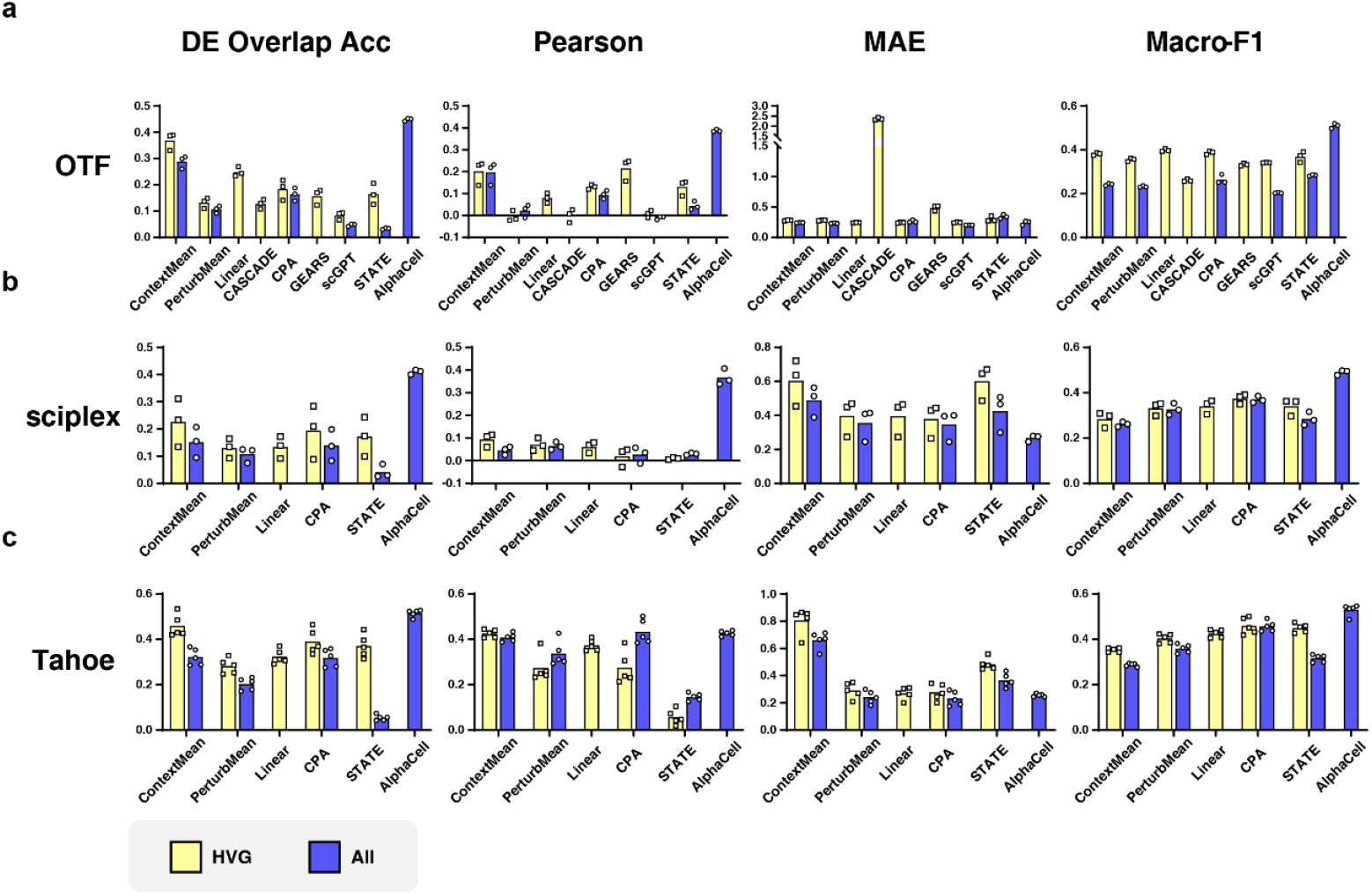
Performance comparison of AlphaCell against state-of-the-art baselines (including STATE, scGPT, GEARS, and linear models) in the compositional generalization task. (a) OTF (transcription factor overexpression; (b), Sciplex (chemical perturbation); (c) Tahoe (large-scale drug perturbation). Metrics evaluate two dimensions of simulation quality: Quantitative Fidelity (Pearson correlation, Mean Absolute Error [MAE]) and Regulatory Logic (DE Overlap Accuracy, Macro-F1). DE Overlap Accuracy measures the successful retrieval of the top differentially expressed genes, while Macro-F1 assesses the correctness of the predicted directionality (up/down-regulation). (HVG——High Variable Gene set; All——Genome-wide gene set)

In terms of quantitative fidelity, AlphaCell demonstrated a superior ability to disentangle perturbation dynamics from cellular context, achieving the highest Pearson correlation with ground truth profiles across all datasets. This performance advantage was most pronounced in the Sciplex dataset (Fig. 4b), where perturbation effects are subtle and notoriously difficult to distinguish from batch noise. While VAE-based approaches and even foundation models like STATE struggled to exceed a Pearson correlation of 0.15, AlphaCell achieved significantly higher fidelity. This divergence validates the robust construction of the Virtual Cell Space by the AlphaCell ***Base Model***: by rectifying the whole transcriptome into a continuous, dense latent manifold rather than relying on noisy HVGs, AlphaCell filters out technical artifacts while preserving the fine-grained topology of the manifold. This allows the AlphaCell ***Flow Model*** to apply perturbation vectors with precise magnitude.

Beyond global correlation, the ability to correctly identify Differentially Expressed Genes (DEGs) distinguishes a mechanistic simulation from mere statistical smoothing. AlphaCell demonstrated a substantial lead in DE Overlap Accuracy and Macro-F1 scores, whereas competitors often produced flat predictions with high precision but limited recall. Notably, expanding baseline models to predict the genome-wise gene set generally resulted in lower DE Overlap Accuracy compared to their HVG-restricted counterparts. This result underscores the critical role of defining perturbations as continuous flows rather than discrete jumps. Traditional models, particularly those reliant on discrete mappings, are prone to overfitting the stochastic noise inherent in individual observations. In contrast, AlphaCell treats the perturbation as a governing law within the Virtual Cell Space. During simulation, the Dynamic Physics Engine drives the cell state embedding along a deterministic trajectory defined by the learned vector field. By capturing the coherent directionality of the perturbation effect across the population, this continuous modeling approach effectively filters out incoherent statistical fluctuations present in raw samples. Consequently, AlphaCell recovers robust biological signals that are otherwise obscured by measurement limitations, distinguishing true regulatory shifts from background noise.

In all, the model’s robust performance across diverse genetic and chemical contexts confirms the effectiveness of the Shared and Routed Mixture-of-Experts (MoE) architecture in mastering complex dynamics. While linear baselines failed to capture the specific directionality of gene regulation (low F1) and discrete mapping models struggled with fine-grained dynamics at the whole-genome scale, AlphaCell successfully transferred learned dynamics to new contexts. This indicates that the Dynamic Physics Engine has learned universal vector fields that are independent of the specific cell state embedding. By successfully decoupling the perturbation force from the cell identity, AlphaCell can apply a learned force to a known initial state in a novel configuration within the Virtual Cell Space, confirming that the model simulates the underlying physical process of state transition rather than simply memorizing cell-perturbation pairs.

### Mechanistic transfer of perturbation dynamics to entirely unseen cellular context

Following the assessment of compositional generalization, we extended the evaluation to cell-type zero-shot generalization —a second level of cellular context generalization ability in the cellular state transition modeling: where the model must predict the response of a cell lineage completely absent from the training data. This regime exposes a fundamental logical inconsistency in traditional HVG-based feature selection strategies. In a zero-shot setting, the transcriptional program of the perturbed state is unknown a priori; thus, it is impossible to predict which genes will exhibit high variance upon stimulation. Relying on HVGs derived from control populations or training datasets assumes that the perturbation response is confined to pre-existing axes of variance, an assumption that fails whenever a perturbation activates quiescent pathways or induces novel cell states. Consequently, all existing HVG-based models operate on an incomplete feature space, rendering them theoretically ill-posed for capturing the full spectrum of a novel perturbation response. In general, most traditional perturbation models rely on learnable, discrete cell-type embeddings, a design choice that structurally precludes them from performing true zero-shot predictions on unobserved lineages without retraining. Only set-based foundation architectures like STATE, which bypass explicit cell-type tokens, are mathematically equipped to attempt this task. AlphaCell circumvents this limitation by operating within the universal Virtual Cell Space constructed by the AlphaCell ***Base Model***. Since this space is defined by the full 19,253-geneset, it provides a mathematically complete substrate that can represent any potential shift in gene expression, regardless of prior variance.

We benchmarked performance against the baseline PerturbMean and the foundation model STATE, across OTF, Sciplex, and Tahoe datasets (Fig. 5). Quantitatively, AlphaCell consistently outperforms baselines, particularly in metrics capturing biological directionality and regulatory precision. When compared to STATE, AlphaCell achieves dramatic, multi-fold performance improvements across all evaluation metrics. For quantitative fidelity, AlphaCell delivers a 2.5-to >10-fold increase in Pearson correlation (e.g., from ∼0.02 to ∼0.2 in OTF, Fig. 5a) and reduces the Mean Absolute Error (MAE) by 30% to 50% across the three datasets. More importantly, for mechanistic accuracy, AlphaCell exhibits a 3-to 6-fold improvement in Differentially Expressed (DE) Overlap Accuracy and a 20% to 50% increase in Macro-F1 scores. The biological significance of these improvements is profound: they demonstrate that AlphaCell goes beyond merely matching the statistical outline of a population; it accurately pinpoints the specific, sparse regulatory genes turning on or off in a completely novel cellular environment. This substantial divergence confirms that AlphaCell does not merely apply a generic average shift, but actively modulates the physical trajectory based on the distinct initial coordinates of the unseen cell type. STATE’s reliance on set-based distribution matching (MMD) fundamentally limits its ability to extrapolate to disjoint manifold regions occupied by novel cell types, leading to near-random correlation and a failure to recall specific DEGs. In contrast, AlphaCell’s continuous flow formulation mathematically applies the generalized vector field to the new anchor point, maintaining robust predictive fidelity even in low-signal regimes like the Sciplex dataset (Fig. 5b).

**Figure 5.**
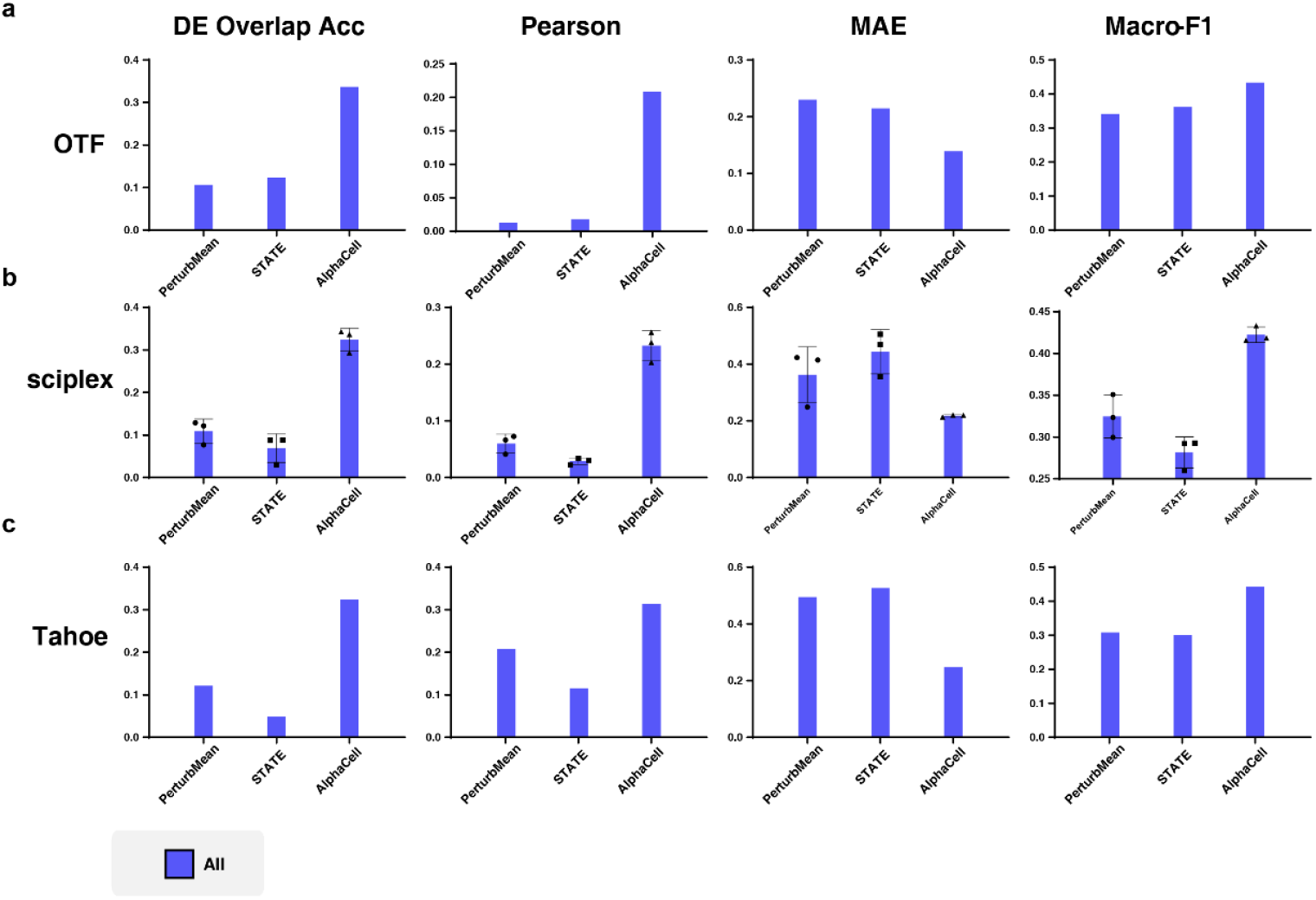
Performance comparison of AlphaCell against state-of-the-art baselines in the cell-type zero-shot task. (a) OTF (transcription factor overexpression; (b), Sciplex (chemical perturbation); (c) Tahoe (large-scale drug perturbation). Metrics evaluate two dimensions of simulation quality: Quantitative Fidelity (Pearson correlation, Mean Absolute Error [MAE]) and Regulatory Logic (DE Overlap Accuracy, Macro-F1). DE Overlap Accuracy measures the successful retrieval of the top differentially expressed genes, while Macro-F1 assesses the correctness of the predicted directionality (up/down-regulation). (All——Genome-wide gene set)

The performance of AlphaCell in this zero-shot setting substantiates the core premise of the Virtual Cell World Model: the universality of the learned state transition dynamics. By training on a comprehensive, batch-corrected manifold, the AlphaCell ***Flow Model*** captures generalized patterns of cellular response that transcend specific training examples. Because the Virtual Cell Space constitutes a continuous and unified substrate, the dynamic laws (vector fields) learned in one region of the manifold can be effectively extrapolated to disjoint regions occupied by novel cell types. The model effectively simulates the physical process of applying a known force to a new object to predict its trajectory. The high fidelity of these predictions indicates that AlphaCell successfully masters transferable laws of cellular dynamics, enabling in silico experimentation on biological contexts that were not empirically tested.

## Discussion

AlphaCell substantiates the paradigm shift toward a generative **Virtual Cell World Model**, systematically validating three theoretical pillars. First, regarding the Virtual Cell Space, our results prove that information compression outperforms feature truncation; processing the 19253-geneset retains critical regulatory drivers that HVG-based methods discard. Second, regarding the Observation Interface, we demonstrate that a massive, knowledge-rich decoder is essential for anchoring abstract latent variables to biological reality, ensuring that mathematical operations within the manifold remain biologically high-fidelity. Third, regarding the Laws of Motion, by replacing discrete mappings with Optimal Transport Conditional Flow Matching, AlphaCell validates that biological dynamics modeled as continuous physical flows is able to capture the non-linear trajectories missed by linear arithmetic.

A critical emergent property of this framework is its ability to distill robust biological signals from technical noise. Single-cell data is inherently plagued by stochastic fluctuations, such as dropout and sequencing depth variations. AlphaCell does not rely on explicit imputation; rather, by constraining predictions to follow coherent vector fields within a smooth latent manifold, the model inherently filters out incoherent statistical fluctuations. This denoised trajectory represents the expected state of the cell population, aligning better with biological ground truth than individual noisy measurements. Furthermore, this manifold smoothness underpins the model’s generalization capability. Because the laws of motion are learned on a unified, batch-corrected Virtual Cell Space, the vector field induced by a specific perturbation remains consistent across the manifold. This allows AlphaCell to mechanistically transfer learned dynamics to the state embeddings of entirely unseen cellular context, effectively simulating how a known force acts upon a novel object.

Despite these advances, AlphaCell represents a foundational step with specific limitations. First, regarding perturbation generalization, the current AlphaCell Flow Model relies on discrete embeddings for perturbation identities. While this enables generalization across cell types, it precludes zero-shot prediction of perturbation. While future updating can be easily incorporate into AlphaCell to achieve zero-shot prediction of perturbation by integrating gene or chemical embeddings to bridge the gap between perturbation space and biological effect. Second, the current virtual world is defined solely by the transcriptome; a complete digital twin must eventually integrate multi-modal layers. Nevertheless, by providing a scalable, high-fidelity simulator for cellular dynamic modeling, AlphaCell establishes a rigorous theoretical framework for digitizing biological dynamics, advancing the transition from descriptive genomics to predictive biological simulation.

## Acknowledgements

This work was supported by the National Key Research and Development Program of China (Grant No. 2025YFC3409300), National Natural Science Foundation of China (Grant No. T2425019, 32341008, 32550745, 82521002, 82541010, U25D9023), New Generation Artificial Intelligence-National Science and Technology Major Project (Grant No. 2025ZD0122802), Shanghai Pilot Program for Basic Research, Shanghai Science and Technology Innovation Action Plan-Key Specialization in Computational Biology, Shanghai Shuguang Scholars Project, Shanghai Excellent Academic Leader Project, Shanghai Municipal Science and Technology Major Project (Grant No. 2021SHZDZX0100) and Fundamental Research Funds for the Central Universities

## Methods

### 1. Data preprocessing process

To ensure that the Virtual Cell Space serves as a mathematically rigorous and biologically faithful substrate, we prioritized strict semantic alignment and quantitative preservation during data preprocessing.

#### 1.1 Bijective entity mapping

A persistent challenge in modeling whole-transcriptome data is the semantic ambiguity inherent in raw genomic annotations (e.g., GENCODE^1^), where a single biological gene symbol may map to multiple Ensembl IDs^2^ due to alternative haplotypes, patch sequences, or pseudoautosomal regions. Such one-to-many mappings introduce structural noise and dilute the gradient during representation learning. To resolve this, we strictly aligned all sequencing data to the HUGO Gene Nomenclature Committee (HGNC) standard^3^. We filtered the input space to a definitive set of 19,253 unique protein-coding genes, establishing a strict bijective (one-to-one) mapping between the biological entities and the model’s input channels. This guarantees that the resulting latent representations encode unambiguous regulatory signals rather than artifacts of sequence alignment.

#### 1.2 Fine-grained expression tokenization

To accurately capture the subtle fluctuations in gene dosage—often critical for identifying low-abundance transcription factors—we employed a high-resolution quantization strategy. Raw Unique Molecular Identifier (UMI) counts were first depth-normalized to counts per 10,000 (CP10k) and log(1+x) transformed. To convert these continuous values into a format suitable for the transformer encoder without sacrificing quantitative fidelity, we multiplied the normalized values by 100 and discretized them into integers:

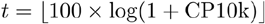

Given that the theoretical maximum of log(1+10^4^) for a cell expressing only a single gene is approximately 9.21, this scaling yields a natural token distribution bounded within a vocabulary size of 1,024. This 100× tokenization strategy preserves expression variations at a resolution of 0.01, capturing minor regulatory shifts while providing a robust, noise-resistant discrete input format^4^.

#### 1.3 Data corpora and scale

The AlphaCell framework was trained across two vast, non-overlapping data regimes. For the construction of the static Virtual Cell Space (Base Model pre-training), we curated an observational corpus of approximately 140 million unpaired single-cell transcriptomes, integrating data from the CZ CELLxGENE Discover census^5^ and the unperturbed baseline samples from the Tahoe dataset^6^. For learning the dynamic laws of motion (Flow Model training), we utilized an interventional corpus comprising over 80 million paired perturbation profiles. This dataset encompasses large-scale genetic and pharmacological screens, including Tahoe-100M, Sci-Plex^7^, and the OTF datasets^8^.

### 2. AlphaCell *Base Model*

The AlphaCell Base Model acts as the Virtual Cell Space builder. Its primary function is to perform Manifold Rectification—transforming the discrete, sparse, and noisy biological observation space (ℝ^19253^) into a continuous, dense, and differentiable Virtual Cell Space that provides the necessary mathematical substrate for dynamic simulation.

#### 2.1 Model structure

Unlike standard autoencoders that maintain symmetric parameter distributions, AlphaCell adopts a highly asymmetric “Inverted Pyramid” architecture, coupling a compact Encoder with a massive, knowledge-rich Decoder.

(1) The Mamba-Transformer Hybrid Encoder (The Latent Manifold Rectifier): The encoder is designed to compress genome-scale data into a strict information bottleneck. It takes the tokenized expression of 19,253 genes as input and utilizes a hybrid architecture comprising 8 alternating blocks of Bi-Directional State Space Models (Bi-Mamba)^9^ and Transformer layers^10^, both augmented with Mixture-of-Experts (MoE) modules^11,12^. Mamba processes linear-time sequence scans to capture global expression trends, while the Transformer provides precise cross-gene attention. Crucially, the encoder utilizes adaptive pooling to forcefully compress the entire transcriptome into 32 continuous latent tokens, forming a 32×128 dimensional latent representation *Z* ∈ ℝ^32×128^. This compression acts as a structural filter: it discards stochastic technical artifacts and forces the network to encode only the essential variables (biological identity and state) into the latent manifold.
(2) The Massive MoE Decoder (The Observation Interface): To ensure that operations within the highly compressed latent space maintain strict biological information, we implemented a 1.2-billion parameter decoder. Serving as the Observation Interface, the decoder consists of 6 Transformer blocks with a wide MoE layer (8 experts, hidden dimension of 2,048) and employs a robust linear projection head to reconstruct the exact log(1+CP10k) expression magnitude (*ŷ*) for all 19,253 genes. This immense capacity ensures that any valid point in the latent space can be translated back into a high-fidelity, whole-genome expression profile, preventing biological hallucinations during dynamic simulation.

#### 2.2 Training stage 1: Base-building

The initial construction of the manifold relies exclusively on the 140 million unpaired observational cells. The objective is to let the model internalize the intrinsic syntax of gene co-expression networks. We employed a Masked Language Modeling (MLM)^13^ paradigm coupled with full-transcriptome reconstruction.

During training, a portion of the input gene tokens ℳ is randomly masked. The total Stage 1 loss is a weighted sum of the MLM loss and the reconstruction loss:

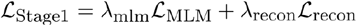

(1) Masked Language Modeling Loss (ℒ_MLM_): A standard cross-entropy loss applied to predict the discretized token ID *ci* for each masked gene *i* ∈ ℳ over the vocabulary size V=1024:

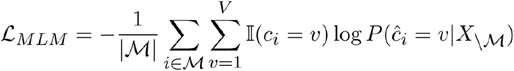
(2) Reconstruction Loss: To reconstruct the unmasked log-normalized expression *y* ∈ ℝ^19253^, the decoder utilizes an L1 loss (Mean Absolute Error). Given the zero-inflated^14^ and highly variable nature of single-cell transcriptomics, the L1 loss is inherently more robust to expression outliers than the Mean Squared Error (MSE). Furthermore, it naturally acts as a sparsity promoter, effectively suppressing technical background noise and preventing the model from hallucinating baseline expressions. For a given cell, the reconstruction loss is computed as:

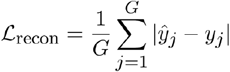

where G=19,253, *y*_*j*_ is the ground truth expression of gene j and *ŷ*_*j*_ is the continuous magnitude predicted by the decoder.

To ensure the resulting Virtual Cell Space reflects the true topological density of biological populations—without allowing highly abundant cell types (e.g., erythrocytes) to monopolize the manifold or rare stem cells to be ignored—we implemented a square-root sampling strategy (τ = 0.5). For a dataset containing N_c_ cells of type, its sampling probability is proportional to N_c_^0.5^. (*τ* = 0.5)This statistical balancing act preserves the core manifold structure dictated by major cell types while securing adequate latent volume for rare transitional states, which are critical for perturbation trajectories.

#### 2.3 Training stage 2: Fine-tuning

While Stage 1 establishes biological semantics, raw single-cell data inherently contains severe batch effects that artificially fracture the latent manifold. Stage 2 is designed to rectify the manifold by stripping away technical artifacts and sharpening cellular identities. However, directly optimizing the latent space for classification often destroys the subtle intra-class variance required to encode perturbation states. To resolve this, AlphaCell employs a Dual-Space Decoupling strategy, defined by the following objective:

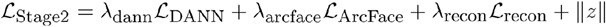

(1) Token-wise Domain Adversarial Neural Network (ℒ_DANN_): To achieve batch invariance, we applied adversarial training^15^. Rather than applying a single global discriminator, we implemented a token-wise DANN that independently penalizes batch signatures on each of the K=32 latent tokens (*z*_*k*_ ∈ ℝ^128^) via a Gradient Reversal Layer (GRL). For *B* datasets (batches):

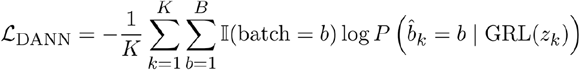 This granular adversarial alignment forces the encoder to thoroughly cleanse batch information from all subspaces, merging identical biological states into a unified coordinate system without destroying perturbation signals.
(2) Semantic Anchoring via ArcFace Regularization (ℒ_ArcFace_): To sharpen cellular identity boundaries without over-compressing the foundational “Physical Space” *Z*, we constructed a separate “Semantic Space.” The 32×128 backbone embedding *Z* is flattened and passed through an independent Multi-Layer Perceptron (MLP) projector to form a Level-2 Semantic Embedding *h* ∈ ℝ^512^. An ArcFace head^16^ is applied exclusively to h. For a cell belonging to class *y*_*i*_ with class weight 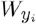, it imposes an additive angular margin *m* on the hypersphere, scaled by *s*:

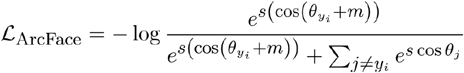

where 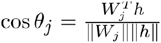. Because the ArcFace gradients are absorbed primarily by the MLP projector, the underlying backbone embedding *Z* remains a continuous, “nebula-like” structure. This decoupling ensures that the backbone retains the essential degrees of freedom required to serve as the mathematical substrate for the subsequent continuous Flow Matching.

### 3. AlphaCell *Flow Model*

The AlphaCell Flow Model functions as the “Dynamic Physics Engine” of the Virtual Cell World. Its sole objective is to learn a deterministic vector field that dictates how a cellular state embedding evolves along a continuous trajectory under a specific perturbation. Because the Base Model has already rectified the discrete biological observations into a continuous, differentiable latent manifold, the Flow Model operates entirely within this foundational substrate.

#### 3.1 Model structure

The Flow Model takes three inputs: the current latent cellular state *z*_*t*_ ∈ℝ^32×128^, the continuous time step *t* ∈[0,1], and the perturbation condition *c*_*pert*_ ∈ 𝒞, which is parameterized via a learnable lookup embedding. To predict the velocity vector field *v* ∈ ℝ^32×128^ while handling the immense mechanistic diversity of distinct perturbations, we engineered a heavily customized Flow Matching Transformer backbone.

(1) Shared and Routed Mixture-of-Experts (MoE): A fundamental bottleneck in training a single monolithic network to simulate thousands of distinct perturbations is catastrophic interference— gradients from functionally opposing perturbations inherently conflict. To resolve this, we replaced the standard dense feed-forward networks (FFNs) in the blocks with an advanced Shared and Routed Mixture-of-Experts architecture. We deployed 16 wide experts (hidden dimension of 2,048), comprising 1 shared expert and 15 routed experts with a top-2 gating mechanism.
(2) Dual Injection of Conditions: To ensure the perturbation signal robustly modulates the cellular trajectory, we implemented a “Dual Injection” strategy. First, for global state modulation, we utilized Adaptive Layer Normalization (AdaLN-Zero)^17^. The perturbation embedding and time embedding are summed and projected to predict the scale (*γ*) and shift (*β*) parameters, directly shifting the distribution of the cell state at every block. Second, for local semantic interaction, we introduced a Joint Attention mechanism where the perturbation token is concatenated with the cell state sequence, allowing the model’s self-attention to explicitly model how the perturbation targets specific latent channels.

#### 3.2 Training scheme

The destructive nature of single-cell RNA sequencing fundamentally precludes the observation of paired temporal trajectories for a single cell (i.e., we cannot measure the same cell before and after perturbation). Consequently, standard supervised regression is impossible. We circumvented this barrier by formulating the task as Optimal Transport Conditional Flow Matching (OT-CFM)^18,19^.

(1) Minibatch Optimal Transport: Instead of relying on static, pre-computed pairings which are computationally prohibitive and prone to overfitting, we applied a dynamic OT strategy[7][8]. During training, within each minibatch, we sample a set of control cells (X_0_) and a set of perturbed cells (X_1_) that share the same biological context and perturbation identity. We then compute the exact optimal transport plan *π*(*x*_0_,*x*_1_)with Sinkhorn algorithm^20^ on-the-fly to pair the cells. This dynamic pairing minimizes the overall transport distance in the latent space, constructing the most biologically parsimonious trajectory (least kinetic energy) from the control to the perturbed population, and acting as a powerful entropic regularizer to prevent the memorization of outlier noise.
(2) Physical Vector Field Matching: Based on the OT pairings, OT-CFM assumes a straight, continuous probability path between the unperturbed and perturbed states:

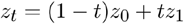

The target velocity of this path is defined simply as the constant displacement:

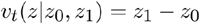

The Flow Model is trained exclusively using a latent physical matching objective, minimizing the Mean Squared Error (MSE) between the predicted velocity and the target displacement.

By relying entirely on this pure latent flow loss, we mandate that the model learns the generalized physical laws of state transitions defined purely by the geometry of the Virtual Cell Space. During inference, we utilize an ordinary differential equation (ODE) solver (e.g., the Euler method) to integrate the predicted velocity field from t=0 to t=1, simulating the continuous biological response to the specified perturbation.

### 4. Task definition

The core objective of the AlphaCell Flow Model is to simulate the evolutionary trajectory of a cellular state under external influence. Formally, given a control cell’s initial state *X*_*ctrl*_ ∈ ℝ^*G*^ and a specific perturbation condition *c* ∈ 𝒞 (e.g., a specific small molecule or gene knockout), the model aims to predict the terminal perturbed state *X*_*pert*_ ∈ ℝ^*G*^. To rigorously verify that the model has learned the underlying physical laws of biological transition rather than merely memorizing statistical correlations from the training distribution, we designed a hierarchical evaluation framework featuring two stringent out-of-distribution (OOD) tasks.

#### 4.1 Compositional generalization task

In this evaluation setting, models are tasked with predicting the outcome of unseen cell-type and perturbation pairings. Specifically, the test set consists of combinations (*Cell*_*A*_, *Pert*_*Y*_) where the cellular identity *Cell*_*A*_ and the perturbation *Pert*_*Y*_ were both independently observed during training (e.g., in the contexts of (*Cell*_*A*_, *Pert*_*X*_) and (*Cell*_*B*_, *Pert*_*Y*_)), but their intersection (*Cell*_*A*_, *Pert*_*Y*_) was strictly held out.

This task evaluates the model’s capacity for compositional generalization^21^. By requiring the model to act as a dynamic matrix completer, it rigorously tests whether the mechanism of a perturbation has been successfully abstracted into a generalized dynamic law capable of reliably operating upon the distinct initial latent coordinates of a newly paired cellular identity. Success in this task indicates that the model has effectively decoupled the perturbation mechanism from the contextual background noise, outperforming traditional linear arithmetic models that tend to entangle the perturbation vector with specific cell-type signatures.

#### 4.2 Cell type zero-shot task

The most extreme test of a world model is its ability to extrapolate physical laws to entirely unmapped territories. In this cell type zero-shot setting, models must simulate the perturbation response of an entirely unseen cell type—a biological context that was deliberately excluded during the training of the Flow Model^22^.

While the specific perturbation condition c has been observed on other cell types, the initial anchor state X_ctrl_ belongs to a novel lineage. This task serves as the ultimate benchmark for the Virtual Cell Space. It evaluates whether the pre-trained latent manifold functions as a truly universal and continuous mathematical substrate, enabling the deterministic vector fields (the “laws of motion”) to be mechanistically transferred and applied to unobserved regions of the biological coordinate system without prior adaptation.

